# CoEVFold suite: user friendly pipelines to visually represent protein coevolution

**DOI:** 10.64898/2026.01.26.701017

**Authors:** Chris L.B. Graham, Liam Cremona, Robin Little, Christopher. D. A. Rodrigues

## Abstract

Multiple sequence alignment (MSA) data underlies current principles in protein folding and protein-protein interaction prediction, from which large language models (LLMs) in tandem with protein datasets, can predict protein structure. However, what is missing are user-friendly tools that enable researchers to predict and demonstrate *coevolution -* the principal input which these MSAs infer. Here we present tools to identify and visualize coevolution, through a pipeline (CoEVFold) that uses basic direct coupling algorithms derived from GREMLIN and alignment of sequences from MMSEQs2. The pipeline generates a visual representation of coevolution for a single protein but can also represent coevolution of homomeric or heteromeric protein complexes, as well as coevolution within protein networks. The input for this pipeline can be an amino acid sequence, or user input protein structures from Alphafold their own files or the PDB database. In validation of CoEVFold’s capabilities, and utilising proteins from known prokaryotic and eukaryotic model systems (*Bacillus subtilis, Escherichia coli* and *Saccharomyces cerevisiae)*, as well as phage proteins, CoEVFold predicts coevolution between proteins known to interact, proteins known to oligomerise, and coevolution in proteins known to be part of a protein complex. Collectively, these suite of tools, named ‘CoEVFold suite’, have broad applicability and provide a useful toolkit to those interested in dissecting protein-protein interactions and networks.

**Availability:** The code is available online at https://colab.research.google.com/drive/1MSSvNTq7KZ4Lr0XTz89vUuK-J3xOTzwS?usp=sharing and Github. https://github.com/MishterBluesky/CoEVFold

**Supplementary information:** Supplementary data is available via Figshare and supplementary materials.

## INTRODUCTION

Coevolution is the evolution of one entity relative to another. It ensures continued function of complex systems, allows for environmental adaptation, and establishes new functions. With respect to amino acids in proteins, coevolution often occurs between two amino acids to enable folding and stabilisation of the protein. It is generally theorised, and has been shown by the success of learning models, that a highly co-evolved amino acid pair within a protein are commonly in physical proximity (1). Thus, coevolution between amino acids within a protein can effectively reveal its 3D landscape, or rank-based coevolution amino acid contact map (1). In addition, coevolution can be used to discover protein self-interactions (homomeric) to assign oligomeric state, or to discover protein-protein interactions between two or more different proteins (heteromeric). The predictive power of coevolution, something which relies on the genetic history of the organism therefore, is great. In the absence of experimental data, it alongside structural tools could be used to solve or work towards solving problems which as of yet are impossible or arduous.

The GREMLIN direct coupling analysis algorithm, established by the Baker lab(1,2) is a powerful coevolution predictor. It was used as the one of the bases towards powerful protein structure prediction tools such as PyRosetta(3); However many coevolution tools have lost support in favour of AI structural prediction tools that give visual representation of the protein tertiary structure. One of the major drawbacks of AI-based tools is the missing information on how proteins folds (secondary and tertiary structure) or protein-protein-protein interactions are derived and modelled. This is important information to understand validity and relatedness of protein structures. This is where coevolution can become an important feature for rationalising modelled protein structures, identifying core structural features and identifying evolutionary relationships.

Coevolution describes the degree to which two molecular elements whether amino acid residues, domains, or entire proteins change in tandem in a predictable way (4,5). Coevolution occurs at the level of species and prophage, between two co-dependent proteins, or two co-dependent amino acids that reinforce a structure. The simplest coevolution to quantify is on the amino acid level; An illustrative example of intramolecular amino acid contact prediction using coevolution is provided in Figure 1A. A sequence alignment of a hypothetical protein from seven species indicates an Arginine 3 to Aspartate 10 pairing (Figure 1) such that when Arginine evolves to Glutamate (E), a reciprocal amino acid change takes place to allow for stabilisation and folding (Figure 1B). If this occurs in the same protein across multiple species, coevolution is likely occurring. The ability to predict coevolution between amino acids, (i.e. likelihood of amino acid Y changing, dependent on amino acid X) can be determined as a coevolution ‘score’ that is calculated by a direct coupling algorithm, which discerns the likelihood of reciprocal change: some amino acids pairs may not always change across different species and therefore harbour lower coevolution, whereas others will change, thus producing a highly coevolving pair. This score can be represented in a 2D grid as a contact map (Figure 1C), and these pairings provide information which build a 3D structure of the protein (Figure 1D). Indeed, until recently some proteins structures were originally solved using coevolution, such as RodA, involved in cell wall remodelling in *Escherichia coli(4)*. Coevolution predictions can also be applied to protein-protein interactions, to provide a interaction network. These are powerful insights for coupling protein sequence to biology, as shown for interactions of RodA with cognate partner protein PBP2(6).

**Figure 1.**
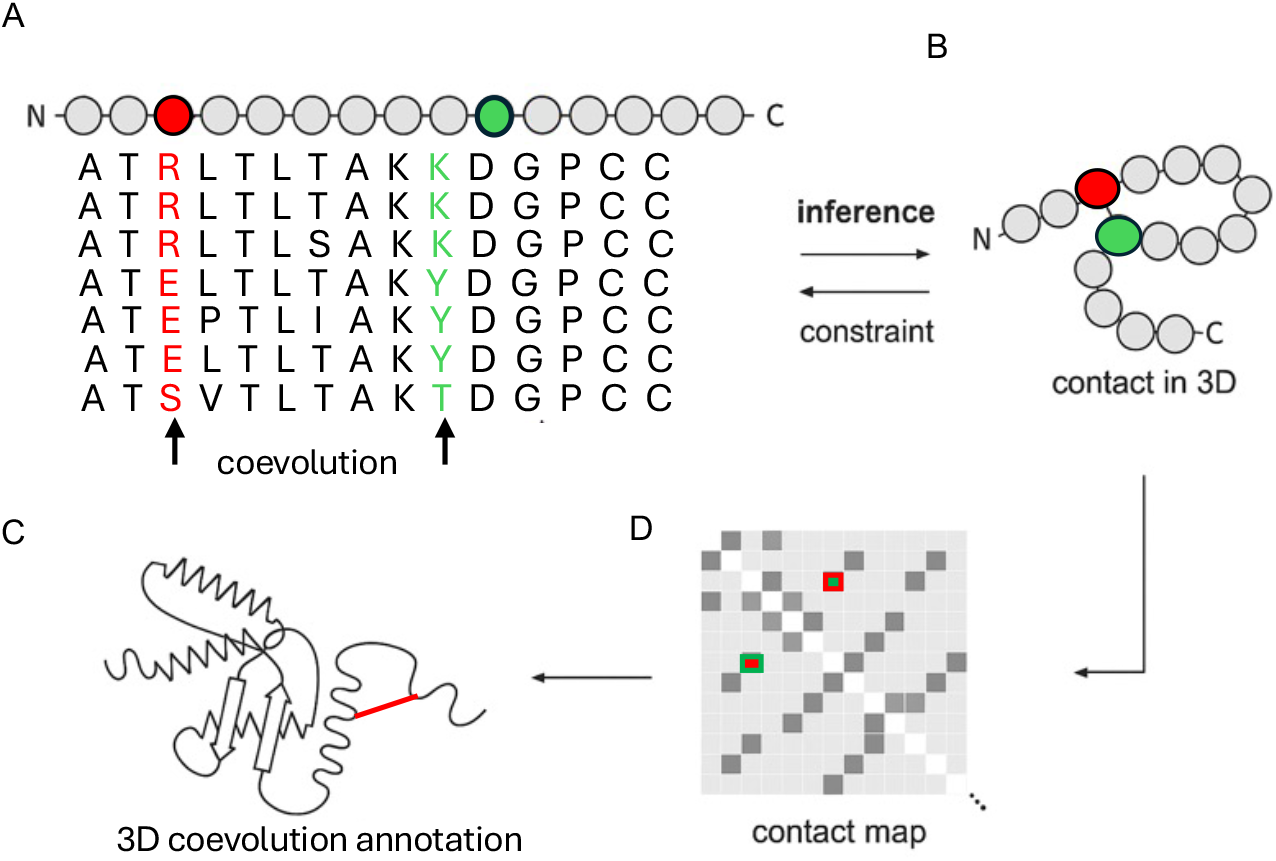
Basic concepts in coevolution. **(A)** Multiple sequence alignment of a protein in seven species, illustrating reciprocal changes in Arginine 3 (R-green) with Lysine 10 (K-green) to ‘stabilise’ a protein **(B)** Contact interface as observed in 2D, if coevolution is strong, to infer a direct physical connection, **(C)** contact map of coevolutionary scores **(D)** Hypothetical overall 3D structure built up of many physical connections informed by coevolution.

Coevolutionary analysis informs on both intramolecular interactions required for folding monomers (Fig 1B), on protomer structure as well as interactions gene networks, and in some cases intermolecular interactions between protomers of a homomeric complex (7). However the differentiation of intermolecular interactions can be difficult to isolate due to protomers often residing in a shared biochemical environment which require parallel changes similar to coevolution coupling (5). Thus, while coevolution can be used as an additional layer of evidence for protein-protein interactions, it is not always representative of a direct intermolecular contact.

Based on the above principles and with this caveat in mind, we have designed a series of python Jupyter notebooks, called ‘CoEVFold suite’ that form an easy-to-use pipeline to study coevolution and use structure to further inform on this coevolution. In these notebooks, we have allowed users to input their own protein structures from protein folding models, or experimentally determined structures, which can be used as a basis for the plotting coevolution between proteins in homomeric or heteromeric states and establishing degrees of coevolution between protein complexes. Finally, we utilise coevolution to determine the likelihood of protein-protein interactions within proteins acting within the same biological pathway.

## RESULTS

We have produced three tools that exploit coevolution for the prediction of protein-protein interactions: 1) CoEVFold - an easy to use coevolution tool; This tool allows users to identify coevolution between two or more distinct proteins of interest; 2) *CoEVFold:4D*, a tool based on CoEVFold to identify coevolution between protomers of homooligomers and 3) CoEVMapper, which predicts protein-protein interactions beyond the protein to itself and a single target creating a network, formed from a target set of proteins within a protein-network.

### Identification of coevolution within two or more distinct proteins in a hetero-oligomer: CoEVFold

Protein complexes underpin many of the catalytic or regulatory functions of cells from across all walks of life. Understanding assembly and interactions as well as the evolution of such complexes is an essential part in discriminating mechanistic function. To identify coevolution between two or more distinct proteins, CoEVfold first relies on MMSEQ2 (8) to automatically an alignment of amino acids based on similarity to a user protein of interest. It then uses direct coupling analysis learning algorithm (GREMLIN)(1), to identify scores for each residue-residue contact and plots this in a grid similar to the illustrations of Figure 1A,(5,7). The protein X to protein Y intermolecular coevolution is then plotted upon a user input predicted multiprotein complex. (9)

Looking at a specific example, of coevolution of bacterial cell envelope protein FtsW with cell division partner penicillin binding protein B (PbpB) (Figure 2A-C) (6,9,10). PbpB homologues have been experimentally shown to interact with FtsW, through their transmembrane helices, and AlphaFold predictions are consistent with this experimental data (6,9,10). However, if a user adds in *other* transmembrane-containing proteins, these will often interact with FtsW at the same site, due to hydrophobic compatibility, thus making pTM scores alone unreliable. Through coevolution mapping with CoEVfold we can generate coevolution scores and map them to the transmembrane region Figure 2C. This provides discrimination between true binding partners and complexes assembled primarily out of charge compatibility. In another example, we plot coevolution of the multiprotein yeast VO type ATPase complex (11). This multiprotein unit must assemble in a very precise manner to enable its dynamic and essential function. The complex shows inter-protein coevolution between chains in its heteromeric state, in particular sharing coevolution with its alpha helical chain encoding genes (Figure 2D-E) such as chain E that forms the central barrel with subunits C, D and to L. This shows distinctive inter-protein coevolution, visible on the coevolution plot (Figure 2D) as well as on the 3D representation (Figure 2E) with interactions indicated by interchain connectors. Each subset of interactions are shown in separate colours.

**Figure 2.**
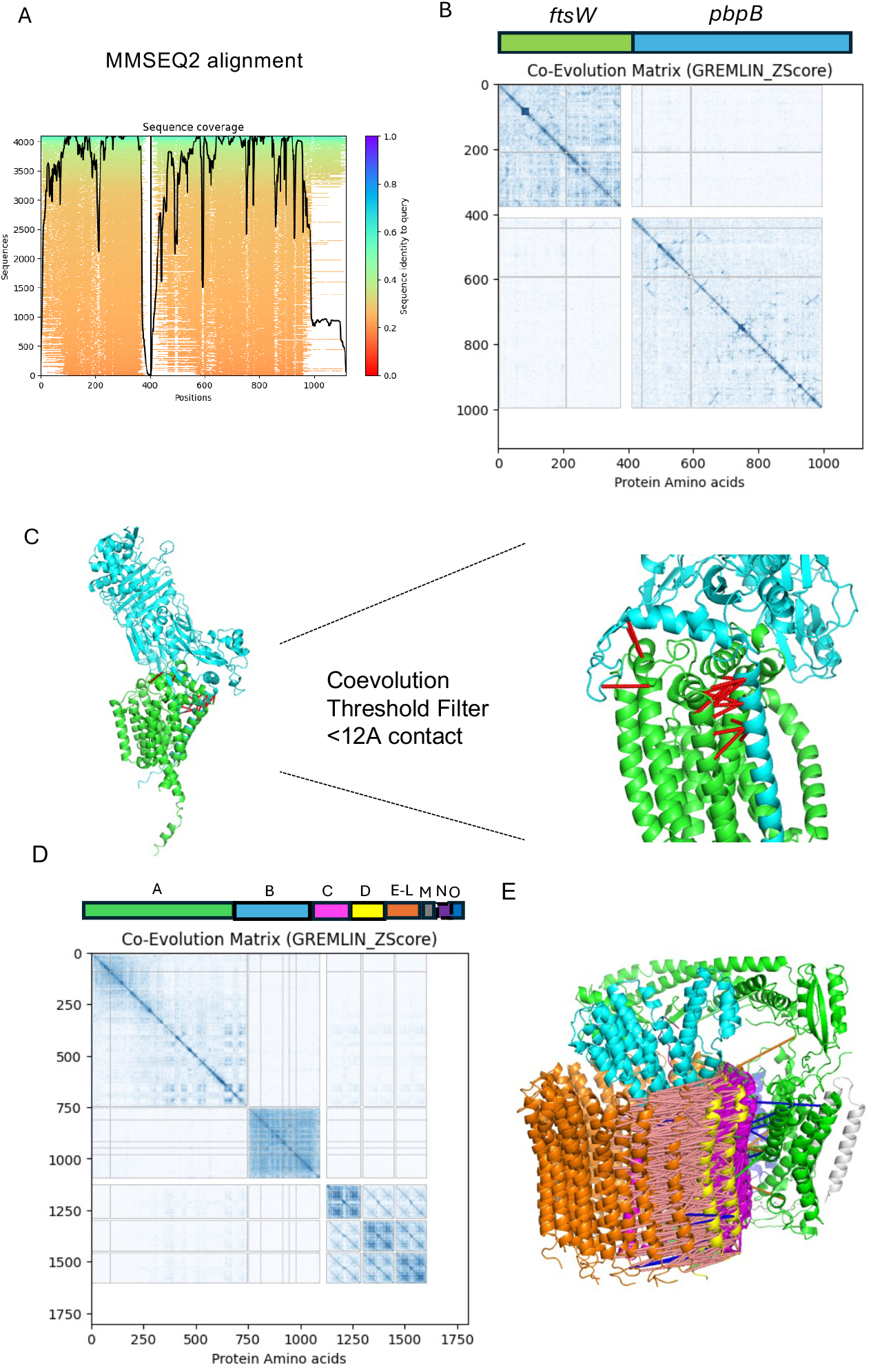
Schematic representation of the CoEVFold pipeline. **(A)** CoEVFold window, MMSEQ2 alignment and a3m production based on query sequence – *B subtilis* FtsW-PbpB >50% sequence coverage. Red – Low coverage/conservation, Blue/Green – High coverage/conservation **(B)** Coevolution of FtsW-PbpB performed using pipeline on transformed alignment. Indicating regions of intermolecular and intramolecular coevolution Dark Blue – coevolution high, White – no observed coevolution Z score min=0, max=3 (**C**) user input structure FtsW-PbpB predicted by Alphafold, inter-protein coevolution has been indicated using red dashed lines = Coevolution at user thresholds (12A, <1.5 Z score). **(D)** Coevolution of query pdb input (PDB ID 7TA0) Cryo-EM structure of yeast VO V-ATPase >50% coverage, Chain indicated above performed using pipeline on transformed alignment indicating regions of intermolecular and intramolecular coevolution Dark Blue – coevolution high, White – no observed coevolution Z score min=0, max=3 **(E)** Query pdb input (PDB ID 7TA0) Cryo-EM structure of Bafilomycin A1 bound to yeast VO V-ATPase Dashed lines represent coevolution links at user thresholds (30A, Z score > 2.5). Each colour represents a unique inter-chain quaternary link subset.

### Identification of coevolution between homo-oligomers: CoEVFold:4D

Protein oligomers are also important in building our understanding of the functionality of cells. In our second tool: CoEVFold:4D, we look for homo-oligomers. Other groups have also used coevolution to achieve this, however, have not provided tools for other groups to do this (3,4), in our tool, we hope to identify coevolution for single proteins in their quaternary, ‘homomeric’ state. CoEVFold:4D works on the idea of uncovering coevolution that isn’t explained by a monomeric structure. By comparing the ‘coevolutionary 2D contact map’ with physical contact maps from the amino acid backbone of respective user inputs from high confidence monomer structures and multimers. These are coevolutionary contacts which are predicted in the 2D map (Figure 3B) but aren’t present in the monomeric state (Figure 3C) are evaluated for attribution to *new inter-protein* links between homomeric forms of the protein (Figure 3E). Dense packing of otherwise unexplained coevolved residues neatly points to a quaternary interaction in that space, thus a tool able to point to these regions in 3D space would allow resolution of homo-oligomer coevolution (Figure 3G-H).

**Figure 3.**
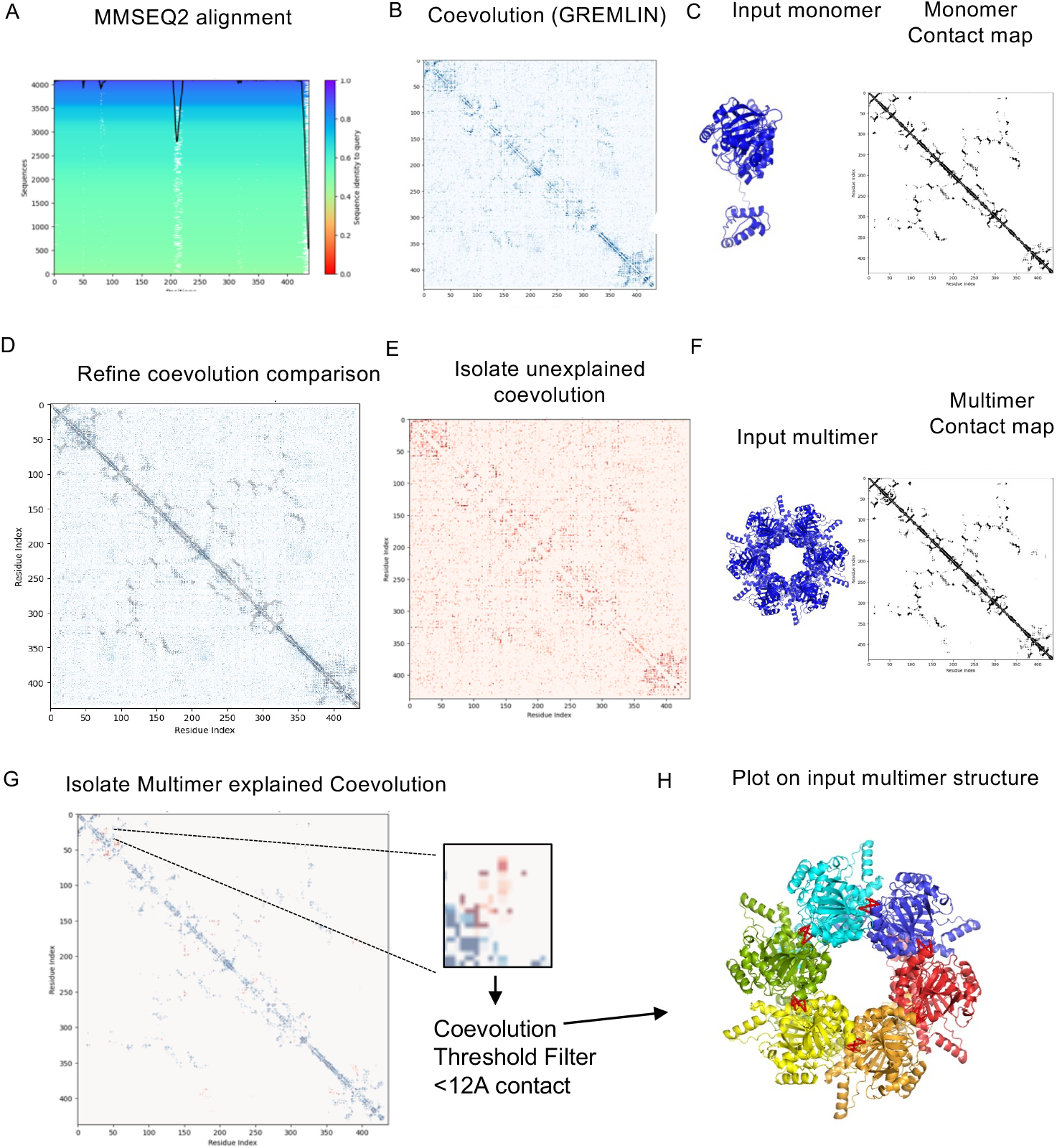
Homomeric inter-protein interaction methodology CoEVFold 4D. **(A)** MMSEQ2 alignment and a3m production based on query sequence – *B subtilis* motor domain SpoIIIE 50%> coverage Red – Low coverage/conservation, Blue/Green – High coverage/conservation **(B)** Coevolution performed using pipeline on transformed alignment. Dark Blue – coevolution high, White – no observed coevolution Z score min=0, max=3 (**C)** User input monomer/lower order multimer and higher order multimer, creating contact map (including intermolecular links). (**D)** Comparison of monomer contact map with coevolution, showing monomer/low order unexplained coevolution (**E)**. (**F/G)** Refinement of search area to coevolution only present in input higher order multimer, red-multimer explained evolution, purple – monomer explained coevolution, (**H)** Optional thresholds of multimer only explained contacts are displayed in a prepared structural .pse file.on input multimer.

This tool is inherently bias and reliant on structural predictions of the monomer. However, it can either be used to confirm an existing predicted multimer or be used to identify regions of contact. We will explore both a guided ‘confirmatory’ quaternary evolution, and an unguided search for unexplained coevolution. There is a caveat here in both modes, coevolution in addition to new interactions could be due to conformational or phylogenetic coevolution noise(11,12). Not all coevolution can be explained by intra-protein and inter-protein interactions (1,5,7). Regions of unexplained coevolution could inadvertently cause users to believe a quaternary interaction is occurring when none is present. These interactions may occur through shared signalling pathways, shared interaction proteins, pH changes, and a conserved local environment, as well as input of aligned sequences. This can be minimised using a combination of coevolution and predicted structures and assemblies. However, these observations are inherently more informed than structural predictions alone.

We applied our tools on a set of multimeric proteins, to confirm that coevolution pairs not explained by monomeric structures can be caused by higher order multimerization of the protein. Using methods as previously described in *B. subtilis* sporulation genes (5). We took an existing known tetramer of *E. coli*, MutS (13), and compared the monomer coevolution with known crystal structure of the dimer. Unexplained coevolution of MutS beyond the dimer was identified in overlaid contact maps (Figure 4 bottom centre). A major cluster of unexplained coevolution was located at the protein C-terminus. *In vitro* mutagenesis studies have indeed confirmed the C-terminus to be involved in oligomerisation (Figure 4) (14). Residues Aspartate 835 and Arginine 840, located at the C-terminus of MutS reduced higher order multimerization (14), correspondingly these residues displayed a high unexplained coevolution score.

**Figure 4.**
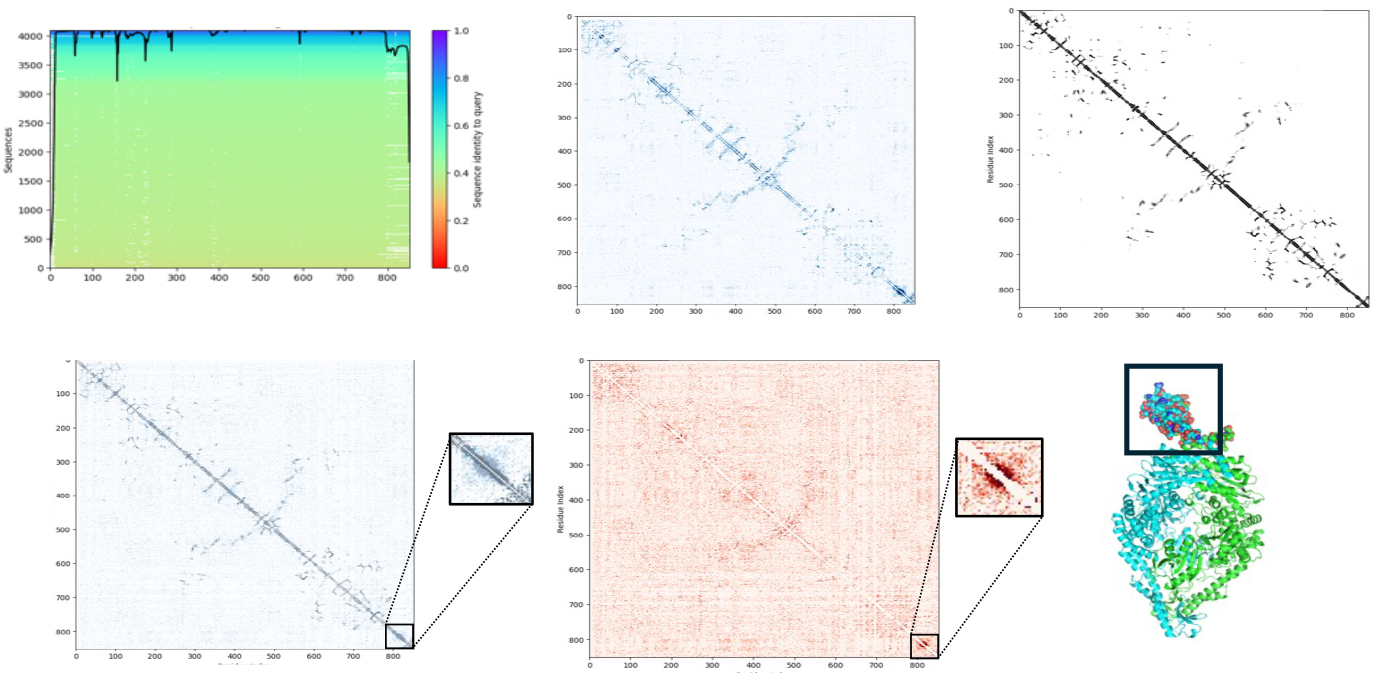
Unexplained Coevolution searching yields similar residues to *in vitro* searches. (top left) MutS MMSEQ2 clustered sequences above 50% coverage Red – Low coverage/conservation, Blue/Green – High coverage/conservation, (top centre) coevolution predicted by tool, Dark Blue – coevolution high, White – no observed coevolution Z score min=0, max=3 (top right) user input dimer 12Å amino acid contact map black (bottom left) Overlay onto coevolution (bottom centre) Unexplained coevolutionary contacts -Red (bottom right) MutS dimer predicted by AlphaFold. Black box, mutated residues shown by in vitro experiments to be responsible for tetramer formation.

We then used this technique to re-identify higher order structures of proteins, in a bias fashion, the second mode of CoEVFold:4D. We tested the known multimers; SpoIIIE’s motor domain (*B. subtilis* inferred by *FtsK* homologue (14), MlaD (*Escherichia coli*)(15) and Tequatrovirus T4 phage tail protein 4UXG(16). Unexplained coevolution contacts unique to contact maps of the multimers were identified (Figure 5). Coevolution interaction clusters that are only present in the multimer, again agreed with known interfaces in structures. with a coevolution cut-off of 2.5 or more deviations above median coevolution (unadjusted Z score) and monomer contact map cut-off threshold of 12Å between backbone residues, thus allowing removal of contacts explained by monomers.

**Figure 5.**
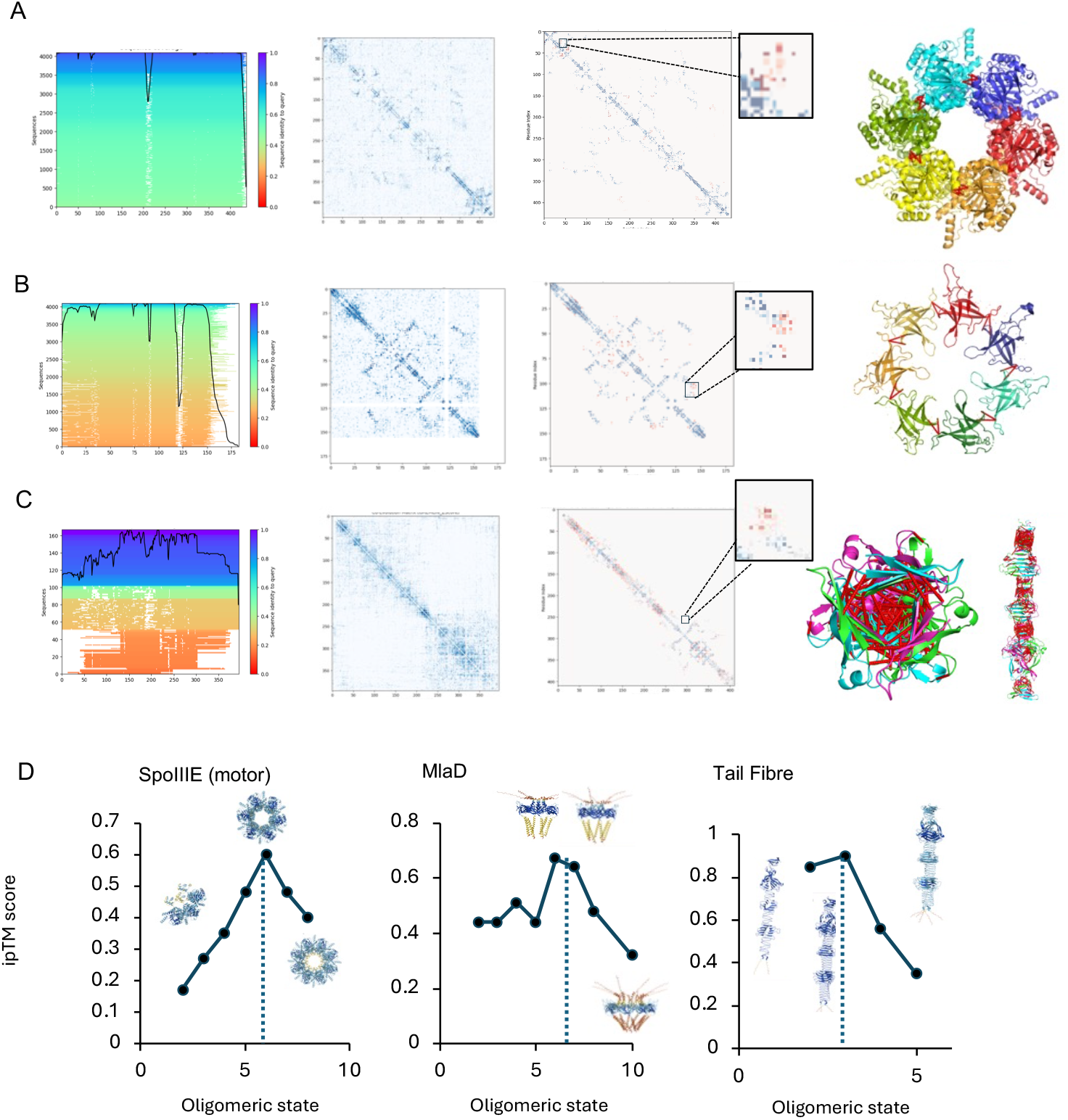
Multimer coevolution and ipTM comparisons, SpoIIIE, MlaD, Tail Fibre. **(A)** SpoIIIE (FtsK homologue) Bacillus subtilis motor domain, (left) MMSEQ2 clustered sequences with more than 50% coverage Red – Low coverage/conservation, Blue/Green – High coverage/conservation, (centre left) Coevolution, Dark Blue – coevolution high, White – no observed coevolution Z score min=0, max=3 (centre right) Multimer only coupling contacts (red), monomer (blue) (right) Contacts in 3D to self between chains above Z score 1 **(B)** MlaD Escherichia coli monomer contrast to crystal PDB ID, **(C)** *Enteroccus* Phage tail PDB ID 4UXG **(D)** AlphaFold3 ipTM score of each protein oligomer at user input oligomeric states.

When comparing contact maps generated from monomeric protein models, small differences in predicted structures at the tertiary level such as backbone bond angles and lengths can create false contacts. Subtly different monomers or an array of monomers in alternative conformations can produce a larger interaction surface for self-contact, thus resulting in unexplained contacts before true ‘quaternary contacts’ can be discerned. To avoid this significant larger clusters indicative of a new protein interface, should be considered first. Therefore, as a final step of inter-molecular interaction selection for biased quaternary contact prediction, coevolution links are plotted in 3D using a generous 12 angstrom or higher distance threshold to eliminate conformer errors. The distance threshold defines the minimum and maximum distance between the backbone of chains in the hypothetical multimer meaning that only interactions between subunits within the threshold are annotated, and not potential differences in tertiary interactions, where interactions may be extremely far apart i.e. above 30 Angstroms Å.

To ensure coevolution analysis is identifying only real interfaces within oligomers ‘Template modelling’ (TM) scores of the inter-protein interactions and in intra-protein interactions; ipTM or pTM respectively should be used as an additional guide, where available. We compared ipTM scores of three proteins at different oligomeric states MlaD, SpoIIE and Phage tail protein of a virus (17), and found that the highest ipTM was observed at the correct multimerization state (Figure 5D), corresponding with coevolution data. MlaD had the highest ipTM in both hexamer and heptameric forms, the tail phage protein as a trimer, and SpoIIIE motor as a heptamer, with coevolution couplings at this interface across model’s indicative of multimerization (highlighted in Figure 5 zoom boxes). The combined use of coevolution with the structural predictions allowed for accurate re-production of existing crystal structures or cryo-EM structures, which cannot be gleaned from a single method alone. This suggests that in the absence of any experimental data both tools need to be combined to accurately predict protein structure and oligomeric states.

In order to investigate oligomer prediction of larger complexes, the combined prediction approach was also applied to *Bacillus subtilis* protein SpoIIIAG (Figure S1) (18). This assembly was not predictable in the state-of-the-art AlphaFold3 using public accounts due to character limits. However, coevolution software is not limited in the same manner. The AlphaFold ipTM suggested the asymmetric hexameric intermediate (Figure S1A) was the best fit oligomeric state for the SpoIIIAG. However, when the coevolution contacts and symmetry restraints are considered the 30-mer becomes the favoured assembly, consistent with experimental data (Figure S1B vs S1D). Thus, highlighting that using coevolution to analyse the interaction surface of these larger assemblies combined with predicted structures of smaller ‘symmetrical units’ provides the most biologically relevant outcome for protein structure prediction.

### Mapping networks of protein interactions using coevolution: CoEVMapper

In addition to modelling protein structures and predicting oligomeric states, coevolution provides a powerful tool for resolving protein interaction networks. Experimentally, it is incredibly challenging to identify a complete network of protein interaction partners within a realistic time scale. Often there are many candidates of potential interactors, and limited genetic phenotypic or biochemical data to validate the interaction, other than co-occurrence of genes. STRING (20,21) provides an excellent resource in gene co-occurrence and neighbourhood which can be applied to discern the likelihood that two genes interact (19,20). However, it does not take into account the product proteins and the role that slight mutations within genes can make in perturbing protein interactions. Further analysis with coevolution data can resolve these features and identify new pathways that are undetected using gene information alone.

A key issue with coevolution is that it is difficult to compare at the scale of genomes due to the computational power required to look at permutations. We wanted to leverage coevolution, similar to how STRING can leverage co-conservation and gene neighbourhoods, to create gene cluster mappings. The idea is simple, we would use the framework of CoEVFold to create an alignment using MMseqs2 (Figure 6A), and a direct coupling contact map (Figure 6B), then rank whole protein inter-protein interactions by the summed coevolution scores. Due to the box sizes and RAM limitations of large contact maps (4000 x 4000 amino acids) which crash typical servers available to users on 12GB RAM python notebooks. We reduced the contact map size and the map was altered to be a representation of a bootstrapped, randomly sampled coevolution between a set of genes (set to 800 by 800 grids). Each square within the grid was representative of a region in the protein. Finally, the coevolution of each protein relative to sampled proteins was compared to respective intra-protein coevolution, to identify potential interaction partners. A ‘high coevolution region’ correction was applied using average product correction (apc), meaning proteins which are intrinsically well conserved or overrepresented in sequences will show to what degree their coevolution is intra and inter protein explained (Figure 6C).

**Figure 6.**
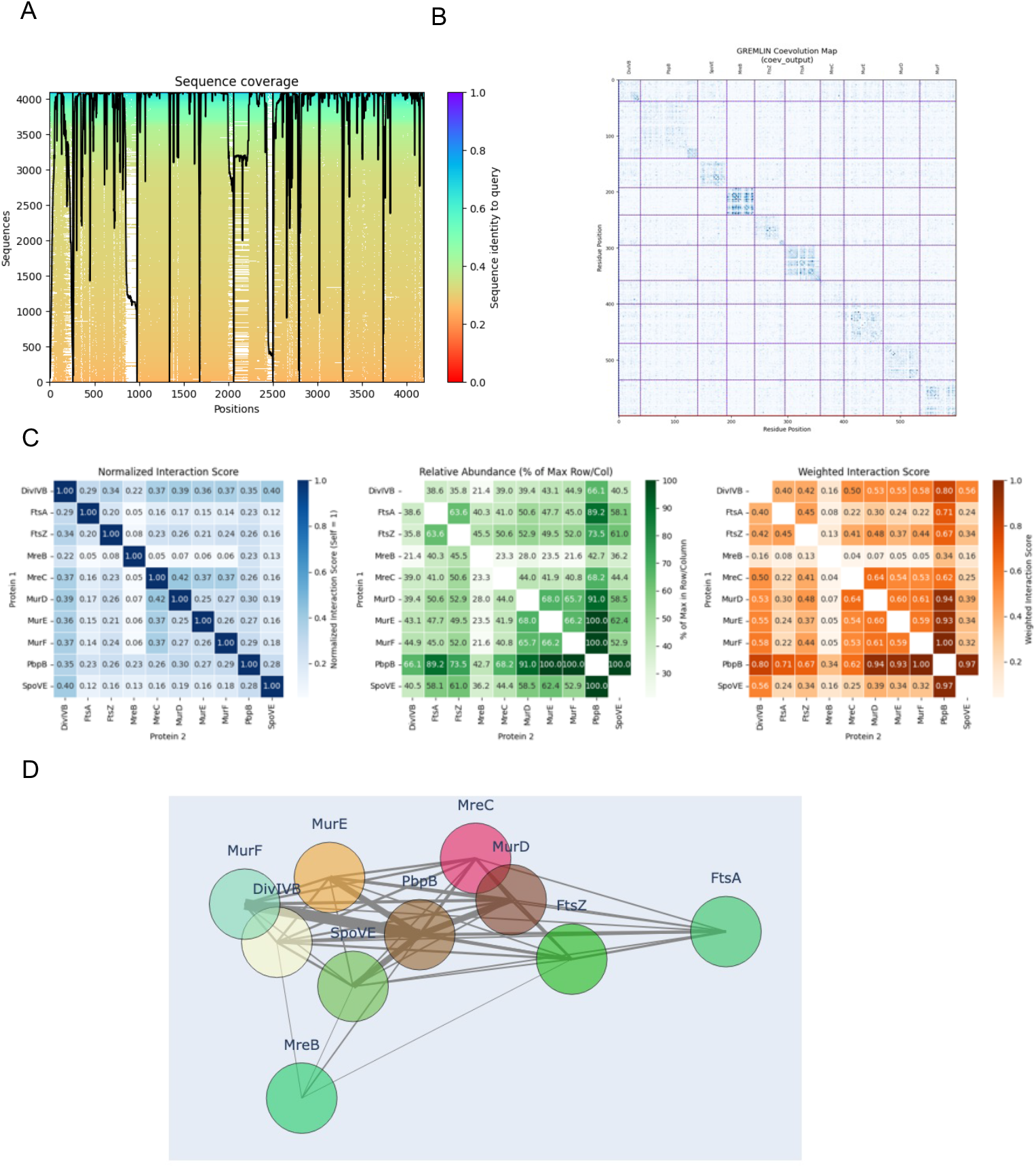
Mapping gene networks using coevolution, CoEVMapper. **(A)** MMSEQ2 alignment of input DCW cluster fasta, Red – Low coverage/conservation, Blue/Green – High coverage/conservation, **(B)** – compressed coevolution (50 bootstraps) Dark Blue – coevolution high, White – no observed coevolution Z score min=0, max=3 **(C)** (left) Normalised interaction score to self (centre) abundance of coevolution above threshold Z score compared to row (right) Weighted interaction score (**D)** – Network of coevolution interactions, line thickness indicating coevolution signal. PbpB coevolution with Mur ligases and SpoVE, FtsA linking with FtsZ.

Our pipeline instigated in CoEVMapper works well for multiprotein systems between 600 and 8000 amino acids in size. Tested examples included the Division and Cell Wall cluster of Bacillus and its closest clades. Currently the system is limited by MMseqs2 inputs as the alignment provider, although this too could be bootstrapped for extremely large queries in the future, <u>or alternatively other filtering methods such as FilterDCA</u>(21). CoEVMapper resolved interactions with peptidoglycan modification genes of *Bacillus subtilis* and its relatives, a system well studied experimentally. We find that DivIB and SpoVE-PbpB interact, FtsA and FtsZ preferably interact, and the MreB Mur-ligase cytoskeletal proteins interact with MreC among all three groups, data congruent with the literature(22). We also analysed a eukaryotic gene cluster (Supplementary Figure 2) using CoEVMapper to compare the coevolution of the RNA polymerase from *Homo sapiens*. Several known core interactions were successfully replicated, particularly those of the well-studied set of proteins (PCF11, CLP1, SSU72, SYMPK, CSTF1, CPSF2, SCAF8 and POL2RA). This highlights that the CoEVMapper method can be applied to study interaction networks across a range of species. Calculating a network however, promiscuity correction could be preferred, as the network interactions can vary greatly dependent on the conservation and therefore level of data in ‘self-coevolution’ a gene has eg: MurE and PbpB are more highly conserved than other proteins. However, this in itself provides information on how some proteins evolve (Figure 6C). This is applied to create a map of hierarchy among coevolution interactions (Figure 6D).

Coevolution mapping of gene networks needs to be done with caution. The process of coevolution refinement of raw data, means that the signal is ‘normalised’ and therefore higher signals are quenched and lower signals in local regions are amplified. This bias can be seen with PbpBs relative propensity to show coevolution interactions with other proteins compared to itself as quite highly scoring, whereas MreB the inverse is true. Bias can be minimized by limiting Z scores to above a user threshold which we have set in the default parameters to above a Z score of 1 or standard deviations above the normal of one and counting the number of interactions and average. Future work will target optimizing these parameters within the program to streamline the network mapping and reducing the risk of user induced bias. Nonetheless coevolution maps from CoEVMapper provide a useful tool for quickly identifying protein interaction networks for hypothesis generation and experimental rationalization. It also provides a platform that streamlines data generation for introduction into other prediction tools such as Alphafold. In our software we provide both weighted and unweighted versions, dependent on the average Z score above the GREMLIN threshold, across lanes. The best approach is not yet clear, and a focus of future work. For example, clade specific use of coevolution and coverage as well as identify cut-offs may be able to get around the biasing of extremely well conserved proteins.

## CONCLUSION

We applied established coevolution prediction methods to generate 2D contact maps of genes of interest. Using a combination of MMseqs2 and the GREMLIN direct coupling analysis algorithm coevolution profiles can be readily generated, providing a powerful tool for inferring biology when combined with protein structure modelling tools or experimental data. There are a variety of applications for coevolution, three of which we have incorporated into pipelines; CoEVFold, CoEVFold:4D and CoEVmapper. These tools effectively map and validate protein interaction interfaces both within homomeric and heteromeric complexes and finally provide a hypothetical gene network visualiser. Our structural interface pipelines in ‘CoEVFold’ predict *in vivo and in vitro* results similar to previous tools, both for homomeric multimerization and heteromeric multimerization. Coevolution is well established in studying protein complexes; however, our goal was to develop a streamlined pipeline that, together with the examples we provide, encourages researchers to explore both heteromeric and homomeric interactions in the context of coevolution and is usable for the unspecialised investigation. By doing so, we aim to reduce the “black box” nature of structural interaction predictions, giving users tools to verify or provide supporting evidence for potential multimeric links within the context of coevolution. Finally, we highlight that coevolution could have unprecedented potential for mapping larger scale gene-cluster wide interactions. Ultimately, we hope that our software grounded in established algorithms will help shed light on how complex structures predicted by new AI tools are established and serve as another means of validating their accuracy, whilst also facilitating new hypothesis generation using genetics as a baseline.

## METHODS

### Prediction of protein structures

All protein structures and multimers listed, unless otherwise stated as an existing pdb structure were created using submissions to the AlphaFold3 server with no modifications. *B subtilis* SpoIIIAG was predicted using residues 40-189. The *B subtilis* SpoIIIE motor domain was predicted using residues 353-789.

### CoEVFold and CoEVFold 4D

CoEVFold works in the following manner. First the user opens a Jupyter notebook explorer in Google Colab, and navigates to CoEVFold, similar to the interface of the widely used ColabFold(23). Then the user uploads a protein structure, or a protein model with two or more proteins interacting in a hypothetical orientation. The query sequences are then derived from the input pdb file and aligned by MMSEQs2 using user defined custom parameters. We recommend a >50% coverage threshold and 95% similarity, as these ensure that the majority of the protein sequences used are similar to the protein of interest and that coevolution is not dominated by a specific clade. After alignment, MMSEQs2 provides an .a3m alignment file of the input amino acid sequence, with the sequence of all amino acids (independent of protein number) conjoined for each species. The conjoined multiple sequence alignment is then subject to direct coupling analysis by a Markov Random Field as originally described by GREMLIN (1). In other words, this coevolution pair algorithm is applied to determine amino acids that co-evolve, and a coevolution score between 0 (no contact) and 1 is determined between the residues in the conjoined sequence. The regions of coevolution can be mapped onto the 3D structure of the protein, with the threshold Z score controlling coevolution displayed. This Z score is based on average intramolecular interactions, as well as intermolecular interactions in the case of heteromers. We recommend based on existing data for inter-protein interactions between heteromers, to use a Z score three deviations above the mean score to reduce false hits. A higher number of sequences input originally will correspond with a higher confidence in the Z scores estimated. For example, consider a protein complex, composed of 4 molecules of protein A and 2 molecules of protein B, obtained by AlphaFold3 or RosettaFold. If coevolution is present at the subunit interfaces, CoEVfold will primarily display coevolution interactions between the chains of each protein in the output .pse file (Figure 2). A higher concentration of high scoring coevolution contacts increases the likelihood of the interaction having occurred over a longer period of evolutionary history. In addition, CoEVFold provides a contact map and a list of the most coevolved residues, which can aid in experimental design such as site-directed mutations to probe oligomeric state.

In CoEVFold, any interprotein interactions which have coevolution and are in contact are included as a connection dependent on the user input coevolution and back-backbone angstrom thresholds. Default parameters; GREMLIN score above 3 and backbone-backbone contact threshold of 15Å. The top-ranking coevolution connections only visible in the multimer are noted and listed as an output, along with their coevolution score, and distance dashes measurements are drawn in PyMOL, to create a pse output of the overall structure explained by coevolution. This shows which amino acids have a high coevolution score and are in contact dependent on user threshold. CoEVFold also notes the reliability. *Eg if the number of sequences provided is less than the total amino acid length, the score is unreliable*. Eg: if 100 sequences are used to find the coevolution of a 300 a.a total length protein complex. CoEVFold is limited by the contact map matrix size, *therefore any interactions which involve complexes in excess of 1000 amino acids may cause the colab to crash, this is a limitation for coevolution discovery. A user could use higher GPU limits possible on A100 GPUs, to assess large complexes such as the V-O type Atpase visualised in Figure 2*.

In CoEVFold 4D, coevolution is scored for each amino acid pair (users are only able to give an input sequence) and users may provide a pdb of the monomer or other lower order multimer (for example from AlphaFold database or a known monomer dimer) and optionally that of the hypothetical higher order multimer, please note: These files must be trimmed to have the same amino acid length as the input sequence. The 12Å carbon backbone to backbone map is derived from the monomeric and multimeric maps. The backbone map is then used as a mask for identification of monomeric and multimeric coevolution interactions. Any residue-residue contacts which have coevolution and are in contact in each structure are included as a connection dependent on the user input coevolution and back-backbone angstrom thresholds. Default parameters GREMLIN score above 3 and backbone backbone contact 15Å. The top-ranking coevolution connections only visible in the multimer are noted and listed as an output, along with their coevolution score, and distance dashes measurements are drawn in PyMOL, to create a .pse output of the overall structure explained by coevolution. This shows which amino acids have a high coevolution score and are in contact dependent on user threshold.

*There are two sub-modes within this homomer search mode on CoEVFold: 4D*.

a. ***Confirmatory quaternary evolution*** – Here the user provides the input sequence, which is aligned using MMseqs2 (Figure 3A) and coevolution is calculated as previously described (Figure 3B). Then the user provides both a monomeric or lower order .pdb file of the protein, as well as a hypothetical higher order multimeric .pdb file predicted by a structural prediction tool such as AlphaFold or TrRosetta (Figure 3C/F). The program finds user thresholded coevolution only explained by the contact maps of the multimer, and plots these on to the structure, saving the result as a PyMOL readable pse file. (Figure 3G). In multimer verification and analysis mode, the contact map of the multimer is also compared (Figure 3H). The coevolution score is plotted on the same contact map for both inputs for plotting graphically, red indicating multimer interactions and blue monomer interactions.
b. ***Unguided quaternary evolution*** – Here the user provides only a low order or monomeric .pdb file of the protein; They can search for unexplained coevolution which might be explained by higher order structures, visualised on a 2D contact map (Figure 3E). The 12Å contact map of the monomeric protein is then used to filter out only coevolution not explained in this 3D map, this gives a set of residues and their scores, which are yet to be explained, which if there is a multimer, especially in areas of high clustering, may be where a multimer forms.

### CoEVMapper

CoEVMapper uses a user input sequence separated by ‘:’ and gene names separated by ‘-’ to establish an input of genes and their respective sequences. It then simplifies these to below a 600.a.a total sequence using random sampling and applies this over a user set number of bootstraps to identify coevolving regions CoEVMapper relies on paired-unpaired coevolution. Finally, the coevolution is compared, using user settings. The code is available on Colab https://colab.research.google.com/drive/1MSSvNTq7KZ4Lr0XTz89vUuK-J3xOTzwS?usp=sharing., and Github; https://github.com/MishterBluesky/CoEVFold/tree/main

## ACKNOWLEDGEMENTS

We would like to thank Sergei Ovchinnikov for his helpful insights into his Github page, and those mentioned in the ‘Beerware license’ of the GREMLIN algorithm. ‘Sergey Ovchinnikov and Peter Koo, as well as Hetu Kamisetty’ who wrote the first GREMLIN algorithm for coevolution searching. We would also like to thank Phillip Stansfeld for his helpful insights and guidance and Melissa Webby for her internal review of the work. This work has been supported by the BBSRC grant BB/X008533/1 awarded to C.D.A.R. We would like to thank the Rodrigues lab for their help in editing the manuscript and testing of the tools, in particular Daniel Dunbar.

