## Supplementary Figure 1 and 2 for "CoEVFold suite: user friendly pipelines to visually represent protein coevolution"

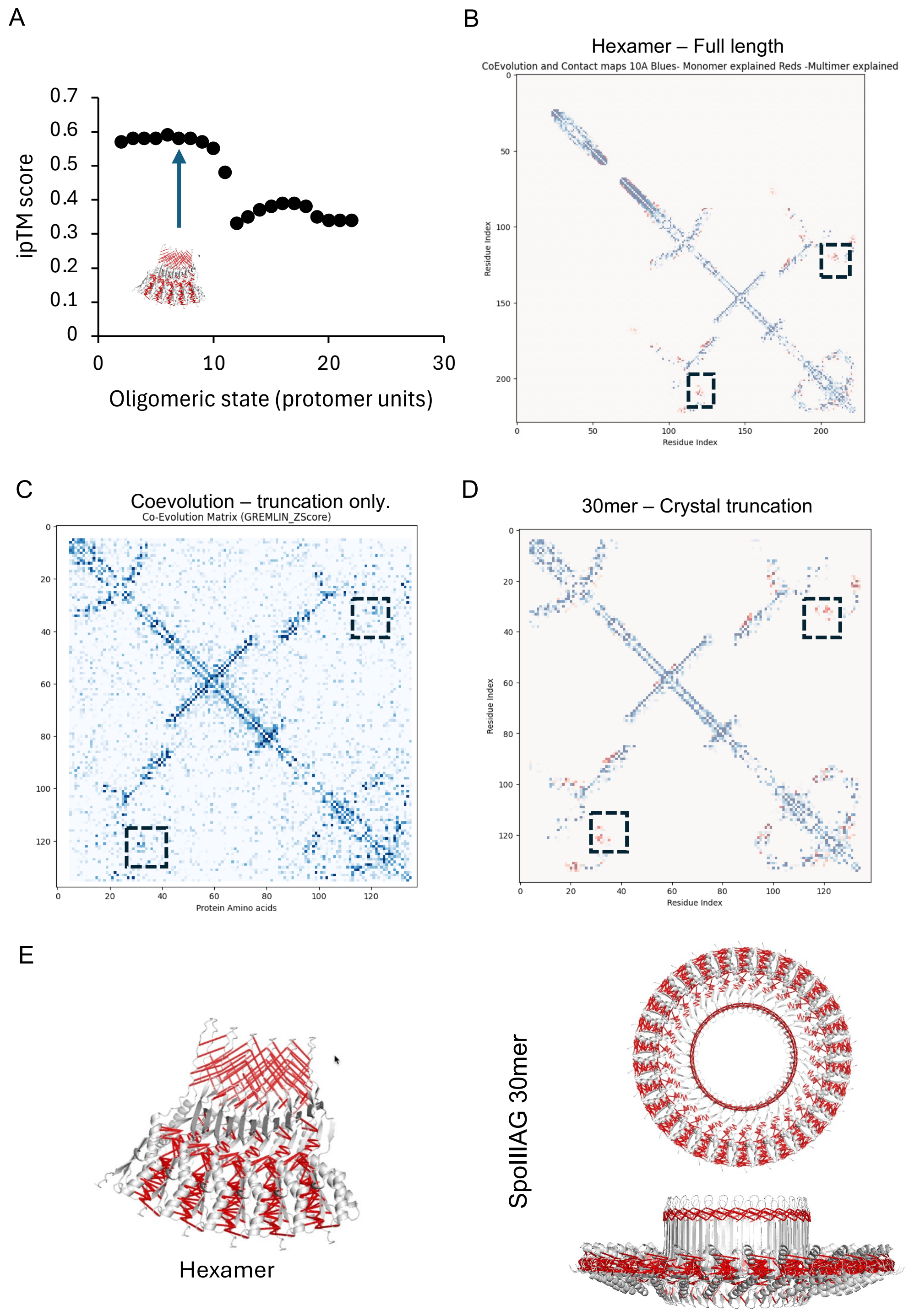


**Figure S1. Existing multimer structures agree with coevolution contacts not found in monomers, but found in best multimer models** A. AlphaFold3 ipTM score of *Bacillus subtilis* SpoIIIAG oligomeric states 2-23. MMSEQ2 sequences B Coevolution of hexamer explained (GREMLIN) multimer only contacts (red) , monomer (blue) and both (red) C Coevolution of SpoIIIAG residues available in pdb structure file. D Coevolution of 30-mer explained (GREMLIN) multimer only contacts (red) , monomer (blue) and both (red). E Quaternary homomer co-evolution interactions on each structure visualized (above Z score 1)

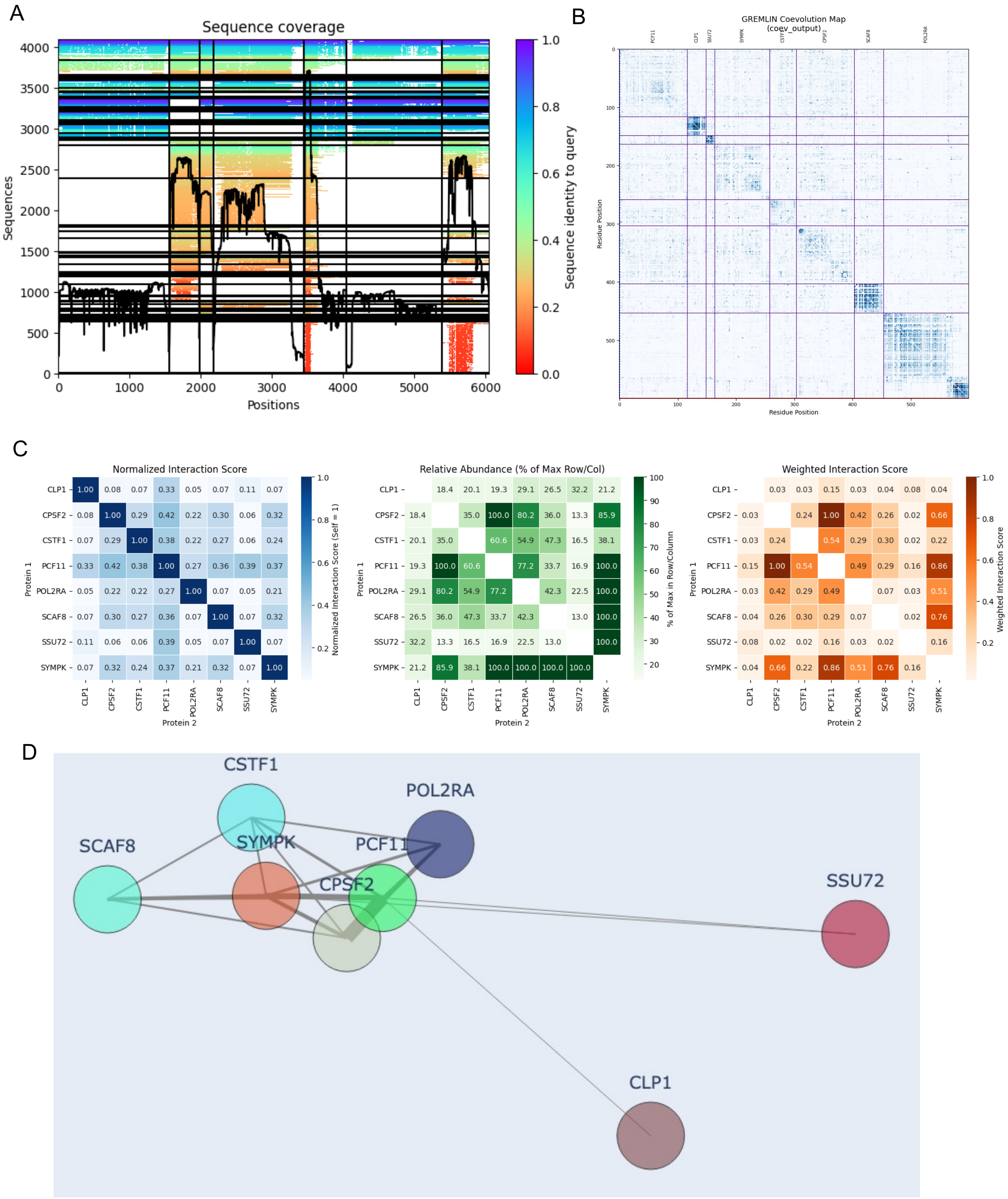


**Figure S2. Mapping human gene networks using coevolution, CoEVMapper**
A MMSEQ2 alignment of input DCW cluster fasta, B – compressed coevolution (10 bootstraps) C – Normalised interaction score to self ii abundance of coevolution compared to row iii Weighted interaction score D – Network of coevolution interactions, line thickness indicating co-evolution signal.
